# Testosterone exposure during fetal masculinization programming window determines the kidney size in adult mice

**DOI:** 10.1101/2025.02.11.637631

**Authors:** Arttu Junnila, Kalle T. Rytkönen, Guillermo Martinez-Nieto, Mats Perk, Hao Li, Jenni Airaksinen, Ida Hyötyläinen, Oliver Mehtovuori, Asta Laiho, Claes Ohlsson, Laura L. Elo, Satu Kuure, Matti Poutanen, Petra Sipilä

## Abstract

Kidney size is sex-dimorphic and regulated by androgens in adult humans and mice. However, the effects of developmental androgen deficiency on kidneys remain elusive. We hypothesized that androgens program future kidney growth during a fetal development. Male mice lacking main testosterone-producing enzyme HSD17B3 had reduced testosterone at embryonic day 15.5, but the concentrations increased by E18.5, creating a short time window of androgen deficiency resulting in reduced kidney size in adult males. In male Hsd17b3-/-kidneys, nephron development was qualitatively normal, but number of glomeruli and proliferation of proximal tubules was reduced. Testosterone supplementation at E14.5-17.5 normalized the renal size in adult Hsd17b3-/-males. Our data suggests that androgen receptor and Hnf4a jointly regulate Igfbp5, FOXO1 and mTOR signaling to convey male-specific kidney growth. In conclusion, we have identified a novel developmental programming effect on male kidneys, where fetal androgen deficiency reduces kidney growth and androgen responsiveness in adult males.

## Introduction

Androgens regulate the development and function of not only reproductive organs, but also e.g. bone, adipose tissue, and kidney. In the kidney, androgens have long been known to regulate gene expression^1,2^, and recent scRNA-seq analyses have further clarified the sex differences^3^. Differential expression of various Na^+^, K^+^, acid–base and organic acid transporters^4^ in female and male kidneys lead to sex differences in kidney functions, such as faster saline load excretion in female rats^5^ and greater urinary ammonia excretion in female mice compared to males^6^. However, the physiological consequences of these sex differences for normal kidney function and in different diseases are largely unknown. Furthermore, male kidneys are larger in both rodents and humans^7^. Orchiectomy and testosterone replacement studies have demonstrated that adult kidney size is controlled by androgens^8,9^, and the effect is mediated via the androgen receptor (AR) located in the proximal tubule^10^. The androgen-induced growth of the kidney in prepubertal/pubertal mice involves cell proliferation first and then cell hypertrophy of proximal tubule cells, and androgens are required to maintain cell size in the adult males^9^. In human kidney transplantation patients, kidney size three months after transplantation depended on the recipient sex, with males having larger tubular area^9^, suggesting that androgens control kidney size also in humans.

We have previously generated a knock-out mouse model for Hydroxysteroid (17β) dehydrogenase 3 (HSD17B3), which is the primary enzyme converting low-active androgen androstenedione (A-dione) to highly active testosterone (T)^11^. The *Hsd17b3*^-/-^ male mice are undermasculinized, with reduced anogenital distance, delayed puberty, and subfertility. They have a similar endocrine imbalance to that of human patients with HSD17B3 deficiency, namely significant testosterone production but a low T/A-dione -ratio. Furthermore, the weights of several androgen-sensitive tissues are reduced, including the kidney, regardless of high T levels in adult mice^12^.

Development of male reproductive organs is dependent on fetal testosterone production, which starts around embryonic day 12.5–13 (E12.5–13) in mice, and reaches a peak around E17–18 before declining again at birth^13^. Inhibition of androgen action during fetal development has been shown to affect the development of the reproductive tract^14,15^. Further work with rats demonstrated a short masculinization programming window at E15.5 to E17.5, which determines the development of the male reproductive tract. Blockage of androgen action only from E15.5 to E17.5 caused underdeveloped internal genitalia, reduced length of phallus and anogenital distance as well as increased number of hypospadias and cryptorchidism^16,17^. Thus, androgens ‘preprogram’ masculinization before actual morphological changes are observed in the developing organs.

Previous work demonstrating the androgen control of kidney growth has been done in post-natal animals. Thus, the role of embryonic T exposure in kidney growth is unknown. In this work we show that androgen action during the masculinization programming window also dictates the size of the kidney in adult mice. The reduced T levels at E15.5 embryos lead to significantly smaller kidneys in male mice, regardless of the high T levels in adult animals. Testosterone treatment of pregnant damns during the masculinization programming window at E14.5 to E17.5 rescued the kidney size to normal in male mice. We further identified signaling pathways putatively mediating the androgen effects in the developing kidney, linking decreased expression of *Igfbp5* to AR and *Hnf4a* regulation and further to downregulation of *Mtor* and *Foxo1*.

## Results

### Kidney size is reduced in adult *Hsd17b3*^-/-^ male mice

One of the androgen-sensitive tissues affected by the lack of HSD17B3 is the kidney, despite the high level of circulating testosterone in adult males^12^. In the current study, the kidneys of 3-month-old *Hsd17b3*^-/-^ male mice were confirmed to be approximately 10% smaller in weight than in WT controls (*P* ≤ 0.01) (Figure 1A). However, histological analysis revealed no morphological abnormalities in the *Hsd17b3*^-/-^ kidneys (Figure 1B).

**Figure 1.**
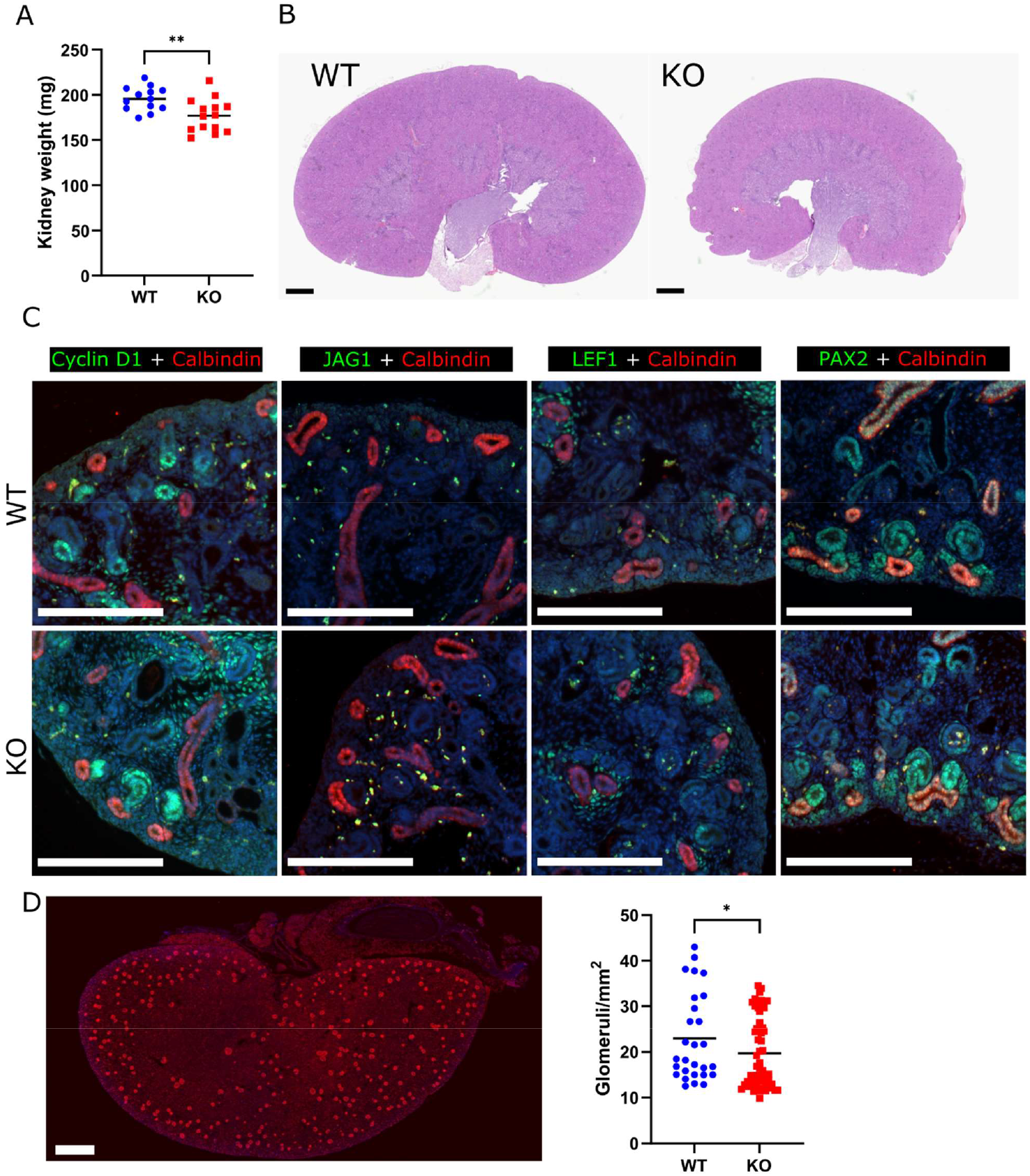
The kidneys of adult *Hsd17b3*^-/-^ male mice are reduced in size but morphologically normal. **(A)** The weight of *Hsd17b3*^-/-^ (KO) male kidneys at the age of 3 months in comparison with WT controls. The lines indicate means. ** = P ≤ 0.01. **(B)** Representative histological images of HE stained WT and KO kidneys. Scale bars: 1000 μm. **(C)** Immunofluorescent stainings of Cyclin D1, JAG1, LEF1 and PAX2 alongside Calbindin in E15.5 kidneys. Scale bars: 200 μm. **(D)** Representative image and quantification of glomeruli at 2 weeks of age via immunofluorescent staining of PODOCALYXIN. Scale bar: 500 μm. The lines indicate means. * = *P* ≤ 0.05

Furthermore, the fetal development of nephron segments and overall kidney patterning appeared normal. We performed immunofluorescent staining of E18.5 kidneys with nephron precursor markers Cyclin D1 (pretubular aggregate to S-shaped body), Jagged 1 (distal renal vesicle, comma- and S-shaped bodies), LEF1 (pretubular aggregate, distal renal vesicle and distal comma- and S-shaped bodies) and PAX2 (nephron progenitors and all nephron structures) along with ureteric bud epithelium marker Calbindin (Figure 1C). These studies revealed no impairment in the induction or patterning of the future nephrons. However, counting the number of glomeruli in 2-week-old male kidneys by staining PODOCALYXIN revealed a 15% decrease in the number of glomeruli per section area in *Hsd17b3*^-/-^ kidney (*P* ≤ 0.05) (Figure 1D).

### The lack of HSD17B3 leads to a delay in fetal androgen production

Although the *Hsd17b3*^-/-^ mice were previously seen to have an undermasculinized phenotype, the circulating testosterone concentrations were normal or increased at the 27-day or 3-month time points, respectively^12^. Therefore, we suspected that the phenotype resulted from an earlier androgen deficiency, during fetal development. Indeed, a clear 5-fold reduction in intratesticular testosterone concentration was observed at E15.5 (*P* ≤ 0.001), with a corresponding 1.3-fold rise in androstenedione concentration (*P* ≤ 0.05) (Figure 2A). However, already by E18.5 the intratesticular testosterone concentration had risen close to normal and androstenedione was at normal levels (Figure 2B). The lack of HSD17B3 activity therefore led to a delay in fetal testosterone production. We have previously shown that by birth, compensatory mechanisms elevate testosterone production in *Hsd17b3*^-/-^ to normal levelThus, androgen deficiency was only observed during a short period in late fetal development, coinciding with the masculinization programming window from E15.5 to E17.5 identified for the reproductive organs.

**Figure 2.**
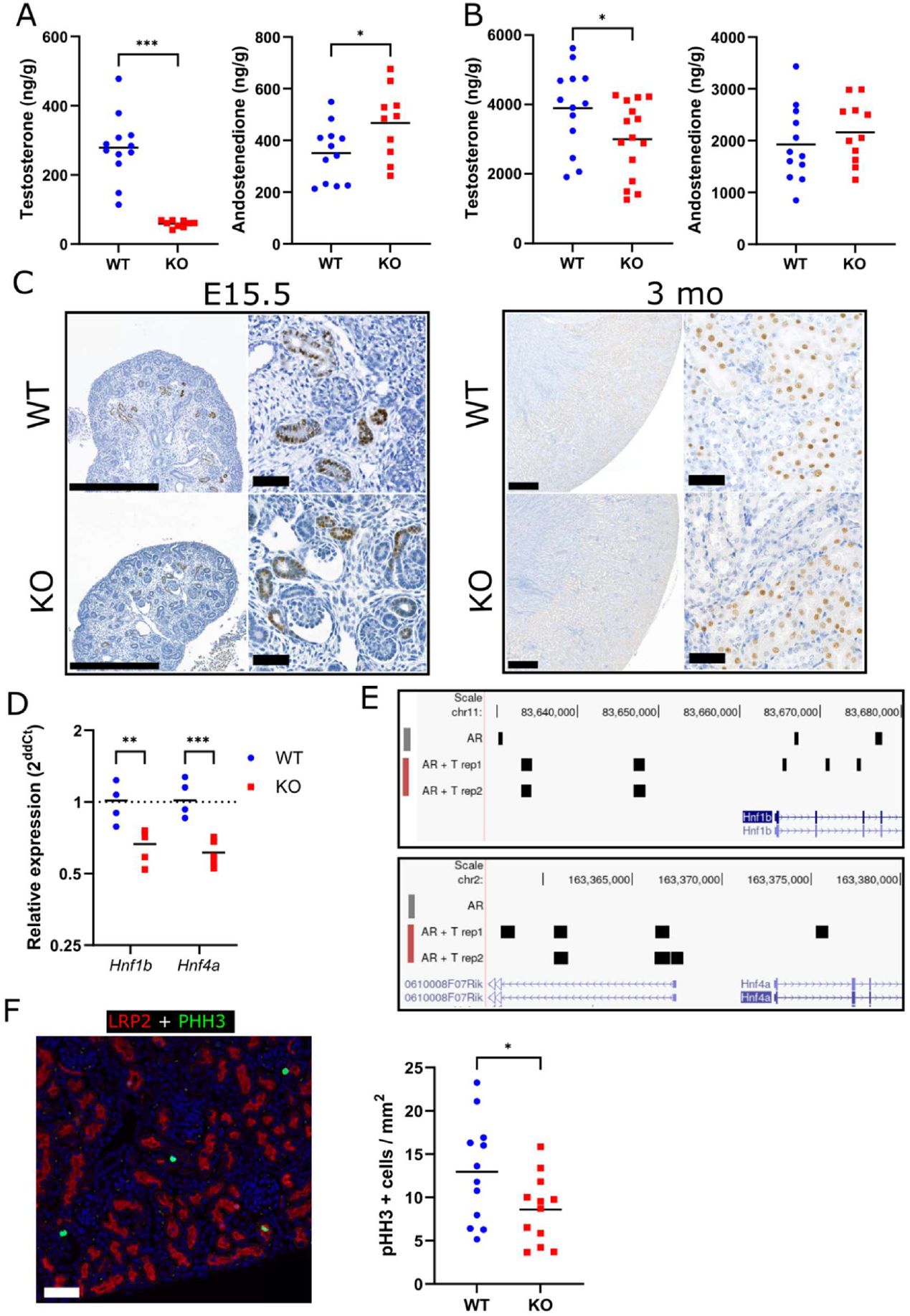
A fetal androgen deficiency in *Hsd17b3*^-/-^ has an effect on androgen-regulated genes and cell proliferation in the kidney. **(A)** Intratesticular testosterone and androstenedione concentrations in WT and *Hsd17b3*^-/-^ (KO) at embryonal day 15.5 (E15.5). The lines indicate means. * = P ≤ 0.05 *** = P ≤ 0.001. **(B)** Intratesticular testosterone and androstenedione concentrations in WT and *Hsd17b3*^-/-^ at E18.5. The lines indicate means. * = P ≤ 0.05. **(C)** Immunohistochemical staining of AR expression in E15.5 and 3-month-old (3 mo) WT and *Hsd17b3*^-/-^ (KO) kidneys. Scale bars on wider field images represent 500 μm, on zoomed images 50 μm. **(D)** Gene expression analysis of *Hnf1b* and *Hnf4a* in E18.5 WT and KO kidneys. The lines indicate means. ** = P ≤ 0.01 *** = P ≤ 0.001. **(E)** Visualization of AR binding peaks near *Hnf1b* and *Hnf4a* in AR ChIP-seq data. **(F)** Representative image and quantification of cell proliferation in 2-week-old kidneys by immunofluorescent stainings of mitosis marker pHH3 (green) and proximal tubule marker LPR2 (red). The lines indicate means. Scale bar: 100 μm. * = *P* ≤ 0.05.

### Fetal androgen deficiency affects gene expression and cell proliferation in the AR-expressing proximal tubule

Expression of AR was observed similarly in WT and *Hsd17b3*^-/-^ male kidneys both during fetal development at E15.5 and in adults at 3 months of age (Figure 2C). At both time points, the expression localized solely in the proximal tubule segment of the nephron, the only region with known sex differences in gene expression^3^.

Hepatocyte nuclear factor family members *Hnf1b* and *Hnf4a* have been linked to the size determination of various tissues, including the kidney^19,20^. The expression of both genes was downregulated in *Hsd17b3*^-/-^ male fetal kidneys at E18.5, after the period of androgen deficiency (Figure 2D). A re-analysis of AR ChIP-seq data from a study by Pihlajamaa *et al*^21^ showed AR peaks both at the promoter and intronic regions of both *Hnf1b* and *Hnf4a* (Figure 2E), suggesting that they are likely directly regulated by AR. Their study also identified *Hnf4a* as a pioneer factor in mouse kidney AR signaling, and the expression of *Hnf4a* was responding to testosterone treatment in adult castrated mice (fold change 1.52, *P* ≤ 0.02)^21^.

To identify the possible changes in the growth of AR-expressing proximal tubules, caused by the reduced androgen action during masculinization programming window, we analyzed cell proliferation in the rapidly growing kidneys of 2-week-old males by double IF staining of low-density lipoprotein receptor-related protein 2 (LRP2) as a marker of proximal tubule segments, and phospho-HISTONE H3 (pHH3) as proliferation marker. The analysis demonstrated that the number of proliferating cells within the proximal tubule area was significantly smaller in *Hsd17b3*^-/-^ kidneys compared to WT (*P* ≤ 0.05) (Figure 2F). The reduced cell proliferation indicates that fetal androgen deficiency does have a persisting effect on the growth of the androgen-sensitive proximal tubules during later development.

### Fetal testosterone supplementation rescues the adult kidney phenotype

To test the effect of fetal androgen deficiency in the development of the kidney, we performed a testosterone treatment of pregnant dams during E14.5-17.5 and analyzed the adult phenotype of the resulting male pups. The fetal testosterone supplementation fully rescued the reduced anogenital distance of *Hsd17b3*^-/-^ males at the age of 3 months (Figure 3A). Interestingly, the androgen supplementation also normalized the *Hsd17b3*^-/-^ kidney weight (Figure 3B). The result demonstrates that fetal androgen exposure does have a role in programming mouse kidney development.

**Figure 3.**
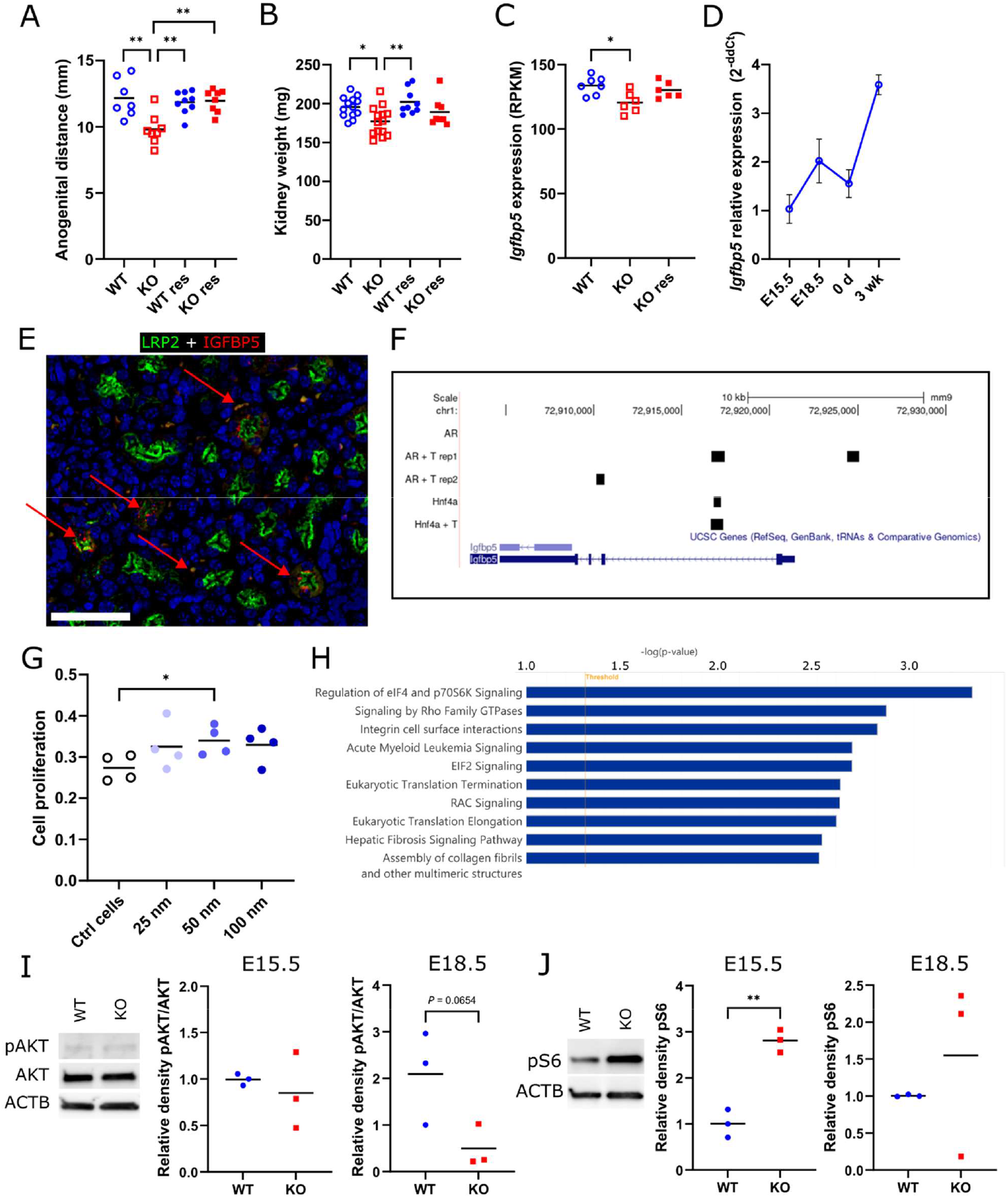
Fetal testosterone supplementation rescues the adult kidney size by normalizing gene expression in the fetal kidney. **(A)** Anogenital distance of 3-month-old KO and WT males without and with testosterone supplementation (res) at E14.5-17.5. The lines indicate means. * = P ≤ 0.05 ** = P ≤ 0.01. **(B)** kidney weights of 3-month-old KO and WT males without and with testosterone supplementation (res) at E14.5-17.5. The lines indicate means. * = P ≤ 0.05 ** = P ≤ 0.01. **(C)** *Igfbp5* expression (RNA-seq) in E16.5 kidneys of WT, KO and treated KO (res) groups (RPKM = Reads Per Kilobase Million). The lines indicate means. * = P ≤ 0.05. **(D)** *Igfbp5* mRNA expression (qPCR) at different age points in WT kidneys. Points indicate means and whiskers SD. d = day, wk = week. **(E)** Immunofluorescent staining of IGFBP5 (red) and LRP2 (green) in 2-week-old WT kidneys. Examples of IGFBP5 staining are marked with red arrows. Scale bar: 100 μm. **(F)** Visualization of AR and HNF4A binding peaks near *Igfbp5* in ChIP-seq data. T = testosterone treatment. **(G)** Proliferation analysis of HEK 239 cells cultured with 0 (Ctrl), 25, 50 or 100 nM recombinant human IGFBP5. The lines indicate means. * = P ≤ 0.05. **(H)** Top 10 most significant canonical pathways obtained from Ingenuity Pathway Analysis (IPA) of DE genes that were normalized in *Hsd17b3*^-/-^ kidney upon fetal androgen supplementation. The top x-axis represents the -log (P-value). **(I)** Representative western blot images in E15.5 and quantification of phosphorylated AKT (pAKT), AKT and loading control β-actin in E15.5 and E18.5 WT and KO kidney homogenates. The lines indicate means. **(J)** Representative western blot images in E15.5 and quantification of phosphorylated S6 ribosomal protein (pS6) and loading control β-actin in E15.5 and E18.5 WT and KO kidney homogenates. The lines indicate means. ** = *P* ≤ 0.01.

To identify genes and pathways affected by fetal androgen deficiency and rescued by testosterone supplementation, we performed RNA-seq analyses of testosterone-treated and untreated kidneys of WT and *Hsd17b3*^-/-^ male pups at E16.5. Gene ontology enrichment analysis of the 416 genes differentially expressed (DE) between untreated *Hsd17b3*^-/-^ and WT kidneys revealed several affected biological processes relevant to the observed phenotype, including those involved in e.g. steroid metabolic process (FDR = 0.0047), regulation of tube diameter (FDR = 0.0047) and tube size (FDR = 0.0050)(Table 1).

**Table 1.** Top 15 gene ontology terms enriched in comparison of vehicle treated WT and *Hsd17b3*^-/-^ fetal male kidneys. FDR = false discovery rate.

| GO ID | Description | FDR |
| --- | --- | --- |
| GO:0015893 | drug transport | 0.00032 |
| GO:0008202 | steroid metabolic process | 0.00472 |
| GO:0008203 | cholesterol metabolic process | 0.00472 |
| GO:0009581 | detection of external stimulus | 0.00472 |
| GO:0009582 | detection of abiotic stimulus | 0.00472 |
| GO:0035296 | regulation of tube diameter | 0.00472 |
| GO:0042730 | fibrinolysis | 0.00472 |
| GO:0050880 | regulation of blood vessel size | 0.00472 |
| GO:0051606 | detection of stimulus | 0.00472 |
| GO:0097746 | regulation of blood vessel diameter | 0.00472 |
| GO:0016125 | sterol metabolic process | 0.00506 |
| GO:0035150 | regulation of tube size | 0.00506 |
| GO:1902652 | secondary alcohol metabolic process | 0.00506 |
| GO:0050982 | detection of mechanical stimulus | 0.01208 |
| GO:0097756 | negative regulation of blood vessel diameter | 0.01365 |

### Androgen deficiency downregulates the expression of *Igfbp5* impacting cell proliferation and signaling

By comparing the DE genes between untreated WT and *Hsd17b3*^-/-^ kidneys, but not DE between untreated WT and testosterone treated *Hsd17b3*^-/-^ kidneys, we identified 292 genes whose expression in *Hsd17b3*^-/-^ was normalized by the testosterone supplementation (Table S1), leading to normalized kidney weight. We compared this set of genes with previously published scRNA-seq datasets from newborn or adult kidneys^3,22,23^ to identify genes co-expressed with AR in the proximal tubules, as possible direct targets of androgen regulation. From the 292 DE genes in *Hsd17b3*^-/-^ kidneys, 11 were found to be expressed in the proximal segment of nephrons (Table S1). One of those, insulin-like growth factor binding protein 5 (*Igfbp5*, Figure 3C), caught our attention as secretory factor that could mediate the effects of androgen action from proximal tubules to other areas of the kidney. In WT kidneys, *Igfbp5* mRNA expression increased by E18.5, then decreased slightly by birth, and rose again at 3 weeks of age (Figure 3D). The proximal tubule localization of IGFBP5 was confirmed by immunofluorescent staining (Figure 3E). Furthermore, reanalysis of an AR ChIP-seq experiment^21^ demonstrated AR binding sites in the promoter as well as first and second introns of *Igfbp5*, one of which also overlapped with a HNF4A binding site (Figure 3F). To establish whether IGFBP5 deficiency could lead to the observed growth defect in *Hsd17b3*^-/-^ kidneys, we treated HEK 293 cells with IGFBP5 recombinant protein. The supplementation of HEK 293 cells with increasing concentrations of IGFBP5 significantly increased their proliferation, as shown by 1.2-fold increment with 50 nM IGFBP5 (*P* ≤ 0.05) (Figure 3G).

We then performed IPA pathway analysis from the 292 genes normalized in *Hsd17b3*^-/-^ kidney by testosterone supplementation. The top ten pathways notably included the regulation of eIF4 signaling pathway, eIF2 signaling pathway and hepatic fibrosis signaling pathway (Figure 3H). All these contained significantly downregulated mechanistic target of rapamycin kinase (*Mtor*) gene (*P* ≤ 0.05) eukaryotic translation initiation factor 4E member 3 (*Eif4e3, P* ≤ 0.05), and forkhead box O1 (*Foxo1, P* ≤ 0.005). Furthermore, IPA network analysis linked *Igfbp5* with FOXO1 and mTOR signaling via AKT function (Figure S1). As AKT activation through phosphorylation is an essential component of these pathways, we analyzed the ratio of pAKT to total AKT in E15.5 and E18.5 kidneys with Western blotting. At E18.5, a lower portion of AKT was phosphorylated in *Hsd17b3*^-/-^ kidneys compared to WT, although the difference did not reach statistical significance due to high variation in WT samples (Figure 3I). mTORC1 activation leads to phosphorylation of ribosomal protein S6, and thus we analysed phosphoS6 from fetal kidneys. However, phosphoS6 was found to be significantly increased in E15.5 *Hsd17b3*^-/-^ kidneys (Figure 3J).

## Discussion

Kidney size in mice and humans is sensitive to androgen exposure, demonstrated by the rapid effect of androgen deprivation and reintroduction on tubule mass and total organ size^9^. Therefore, it was not necessarily surprising that a mouse model lacking a crucial enzyme of testosterone production would have reduced kidney size. However, the fact that the adult *Hsd17b3*^-/-^ male mice still produced high amounts of testosterone suggested a mechanism regulating the growth potential of male kidneys independently from androgen exposure in adulthood.

In this study, we observed that the lack of HSD17B3 in male mice led to testosterone deficiency only during a short fetal time window. The size difference of the kidneys, however, mostly develops after puberty^9^. It has previously been shown that the growth potential of the male reproductive organs in rodents is determined by androgens during the fetal masculinization programming window^16,17^. The target size is reached through later androgen exposure but the final organ size will remain reduced if the early exposure is insufficient.

We have now shown that a similar programming window exists for the mouse kidney: the smaller organ size in adult *Hsd17b3*^-/-^ male mice results from a fetal testosterone deficiency. High adult testosterone level is unable to rescue the kidney size, which is however normalized by fetal testosterone supplementation within the identified developmental time window. This is also supported by our previous results on another mouse model lacking both HSD17B1 and HSD17B3 activity. There we demonstrated a more complete abolishment of fetal testosterone production, resulting in an even greater reduction in adult kidney weight, yet the serum steroid profile after puberty was identical to *Hsd17b3*^-/-^, including high testosterone^18^.

The proximal tubule is the only structure of the kidney known to express AR and respond to androgens^3,9,10^. Here, too, we detected AR expression only in the cells of the proximal tubule, beginning from the fetal period. Moreover, significant down-regulation of *Hnf1b* and *Hnf4a* expression was observed in E18.5 kidneys of *Hsd17b3*^-/-^ males. As both of those genes were shown to be androgen regulated^21^ and ChIP-seq data showed AR binding to promoter and intronic regions of both genes, the observed down-regulation in our study could be due the lack of androgens during preceding days in *Hsd17b3*^-/-^ kidneys. Interestingly, HNF4A has been shown to regulate differentiation of proximal tubule progenitor cells into mature proximal tubule cells^19,24^. Thus, the detected down-regulation of *Hnf4a* could putatively lead to a smaller number of proximal tubule cells and subsequent defect in proximal tubule cell proliferation at 2 weeks of age and hence smaller kidneys in *Hsd17b3*^-/-^ males. As *Hnf4a* was recently predicted to regulate male-biased transcriptome in the kidney, having considerable overlap in target genes with AR^25^, and identified as pioneer factor for AR in the kidney^21^, the growth of male kidney would likely be mediated through their co-operative control of gene expression in the male proximal tubule in response to androgens.

In our RNA-seq analysis, a relatively small number of genes, 292, were significantly changed between the genotypes in E16.5 embryonic kidneys but normalized by testosterone supplementation. From those 292 genes, only 11 were found to be expressed in the proximal tubules of newborn kidneys^3,22,23^, making them possible targets of direct AR regulation. We focused on *Igfbp5* that has been previously reported to be among androgen-regulated genes in the adult mouse kidney^21^. We demonstrated that the gene has associated AR binding sites that also overlap with HNF4A binding, suggesting that IGFBP5 expression is one of the factors in male proximal tubule development co-regulated by both AR and HNF4A^25^.

IGFPB5 is a secreted factor that conveys its effects either via regulating IGF functions or independently of IGFs^26^. The lack of *Igfbp5* has been linked to e.g. increased body size in knock-out studies, likely due to increased IGF-1 activity in the animals^27^. On the other hand, IGFBP5 has been shown to increase the proliferation of papillary thyroid carcinoma cells^28^ and mouse osteoblasts independently of IGF-1^29^. Here too, we demonstrated that the addition of recombinant IGFBP5 increased the proliferation of cultured human embryonic kidney (HEK 293) cells. Therefore, it is possible that it has a similar direct effect on the kidney in vivo.

Studies in human prostate cancer cell lines suggest that overexpression of IGFBP5 decreases PI3K and pAKT levels^30^, and regardless of whether the effect is IGF-1 dependent or independent, it would likely affect mTOR signaling. Our IPA network analysis linked downregulated *Igfbp5* with downregulated *Foxo1* and *Mtor* via predicted downregulation of AKT activation, and indeed, we observed reduced pAKT levels in E18.5 kidney of *Hsd17b3*^-/-^ males. IGFBP5 also contains a functional nuclear localization sequence^26^, and a recent study with rat primary hypothalamic cells suggested that IGFBP5 increased the protein levels of MTOR and AKT^31^. Thus, the observed downregulation of *Foxo1* and *Mtor* in this study could be caused by direct transcriptional regulation by IGFBP5. Moreover, *Mtor* itself has previously been identified as both a male proximal tubule specific and androgen regulated gene^3,21^. Nevertheless, the downregulation of *Igfbp5* and *Mtor*, alongside reduced pAKT levels in *Hsd17b3*^-/-^ kidneys, did not reduce downstream phosphoS6 levels in E15.5 and E18.5 kidneys, with a significant increase instead observed at E15.5. Our findings are, however, in line with previous studies. Lower *Mtor* expression has been shown to affect kidney size and reduce nephron number^32^ as also seen in our study. On the other hand, in adult male mice, orchiectomy increases phosphoS6 levels^9^ similarly to T deficient E15.5 *Hsd17b3*^-/-^ kidneys. Importantly, S6 phosphorylation can also be the result of e.g. the Ras-ERK pathway^33^ and thus does not necessarily reflect the mTOR activity. These identified changes in the interconnected signaling pathways affecting cell growth, proliferation, and viability are all likely to contribute to the observed kidney growth impairment, but further studies are needed to clarify the causality of observed changes.

Our studies give new insights into human HSD17B3 deficiency, which causes a disorder of sex development in XY individuals. The results have relevance for patients whose endocrine phenotype is very similar: fetal androgen deficiency, followed by virilization after puberty as testosterone production is activated^12,34^. There is evidence that a programming window for reproductive tissues may also exist in humans, theorized to be between 8-14 weeks of gestation, and adult mouse and human kidneys react to androgens in a very similar way^9,16,35^. Thus, a similar mechanism of kidney developmental programming in humans is plausible. Reduced kidney size comes with a diminished nephron number, which increases the risks of chronic kidney disease, hypertension and cardiovascular problems^36^. Importantly, the findings are relevant not only for individuals with HSD17B3 deficiency but also for other disorders such as 5α-reductase deficiency, androgen insensitivity syndrome, or environmental exposure to hormonal disruptors. However, further work is needed to clarify the possible consequences of the changes in kidney development. The sex differences in the kidney can be assumed to have a physiological function. Sex-atypical kidney phenotype can then have effects on their capacity to perform their function, if not in normal conditions, then during infections, or challenges such as high sodium diet or heavy use of e.g. certain painkillers^37,38^.

In conclusion, we showed that the lack of HSD17B3 and the resulting disruption of testosterone production leads to reduced kidney size in *Hsd17b3*^-/-^ male mice. The origin of the effect is in a brief fetal time window of androgen deficiency at E15.5-E18.5. This results in developmental changes that cannot be reversed by higher testosterone later in development, but are rescued by fetal testosterone supplementation. One likely mechanism behind the effect is a decrease in AR- and HNF4A-regulated expression of IGFBP5 in the proximal tubule area, leading to decreased proliferative potential in the proximal tubules.

## Methods

### Mouse model

The generation and genotyping of the *Hsd17b3*^-/-^ mouse line has been described by us previously^12^. Mice were housed under a controlled environment (12h light cycle, temperature 21 ± 3°C, humidity 55% ± 15%, specific pathogen-free) at the Central Animal Laboratory of the University of Turku. Soy-free SDS-RM3 chow (Special Diets Service) and tap water were available ad libitum. All animal experiments were approved by the Finnish Animal Ethics Committee (licenses no. ESAVI/7487/04.10.07/2013, ESAVI/41729/2019, and ESAVI/23322/2023) and fully met the requirements as defined by the U.S. National Institutes of Health guidelines on animal experimentation.

### Measurements and tissue sampling

The anogenital distance was determined by measuring the distance between the external genitalia and the anus with a digital caliper (Hogetex). The measurements were done from neonatal animals and at the age of 3 months (WT n = 7, *Hsd17b3*^-/-^ n = 8).

For the collection of tissues, mice were sacrificed at predetermined ages by carbon dioxide asphyxiation, followed by blood collection via cardiac puncture and cervical dislocation. Kidneys were weighed and snap-frozen in liquid nitrogen or fixed for histological analyses. For samples from fetal time points, pregnant dams were sacrificed after timed matings at 15.5 days or 18.5 days post coitum (dpc) and kidneys and testes were collected from embryos.

### Testosterone rescue experiments

On 14.5 – 17.5 dpc, pregnant dams were treated with testosterone propionate (Sigma-Aldrich) in corn oil (Sigma-Aldrich), administered s.c. once daily at 20 mg/kg. Male *Hsd17b3*^-/-^ (n = 8) and WT (n = 9) pups were left to mature and were sacrificed at the age of 3 months.

For fetal kidney RNAseq, pregnant dams were treated either with testosterone propionate or with corn oil as vehicle control, and sacrificed at 16.5 dpc, 2 hours after the third testosterone propionate injection. The embryos (vehicle-treated WT n = 4, vehicle-treated *Hsd17b3*^-/-^ n = 6, T-treated WT n = 4, T-treated *Hsd17b3*^-/-^ n = 6) were collected and the kidneys were snap-frozen in liquid nitrogen for RNA extraction.

### Steroid analyses

The testes were homogenized in sterile deionized water 1:10 (w/v) using an Ultra-Turrax homogenizer (IKA-Werke). The concentrations of androstenedione and testosterone in the testes of E15.5 (WT n = 12, *Hsd17b3*^-/-^ n = 10) and E18.5 (WT n = 13, *Hsd17b3*^-/-^ n = 15) fetuses were analyzed by a validated GC-MS/MS, with the quantification limits of 12 pg/ml and 8 pg/ml respectively, as previously described^39^.

### Histological analyses

Kidneys from 3-month-old mice were fixed in 10% neutral buffered formalin (FF-Chemicals) for approximately 24 hours at RT. The samples were then dehydrated, embedded in paraffin, 5 μm sections were prepared, deparaffinized and rehydrated in a xylene-ethanol series, stained with hematoxylin and eosin (HE), and analysed via light microscopy.

### Immunohistochemistry and immunofluorescence staining of tissue sections

The localization of the androgen receptor in WT mouse kidneys at E15.5 and 3 months of age was visualized with immunohistochemistry. Antibodies are listed in Table 2. The antigen retrieval for tissue sections was performed in a pressure cooker in 10 mM citrate buffer. Blocking was done with 3% bovine serum albumin (BSA) in PBST (PBS with 0.05% Tween). The primary antibodies were diluted in the blocking solution and incubated at 4°C overnight. Endogenous peroxidase activity was blocked with a 20-minute wash in 1% H2O2. The color formation was achieved with Dako EnVision HRP Labelled Polymer Anti-Rabbit and Dako liquid DAB+ (K4003; Agilent) followed by a brief background staining with Mayer’s hematoxylin and mounting. The results were analyzed via light microscopy.

**Table 2.** Primary and secondary antibodies and their use. IH = immunohistochemistry, IF = immunofluorescence, WB = western blot.

| Antibody | Species | Manufacturer | Cat # | Dilution | Used in |
| --- | --- | --- | --- | --- | --- |
| anti-AR antibody | Rabbit | Abcam | ab133273 | 1:500 | IH |
| anti-Cyclin D1 | Rabbit | Thermo Scientific | RM-9104-S1 | 1:500 | IF |
| anti-Jagged 1 | Rat | Hybridoma Bank | TS1-15H | 1:300 | IF |
| anti-LEF1 | Rabbit | Cell Signaling Technology | 2230 | 1:400 | IF |
| anti-PAX2 | Rabbit | Invitrogen | 71-6000 | 1:400 | IF |
| anti-Calbindin | Goat | Santa Cruz Biotechnology | sc-7691 | 1:500 | IF |
| anti-Podocalyxin | Goat | R&D Systems | AF1556 | 1:400 | IF |
| anti-Phosphohistone H3 | Mouse | Cell Signaling Technology | 9706 | 1:200 | IF |
| anti-Lrp2 | Rabbit | Abcam | ab76969 | 1:1000 | IF |
| anti-IGFBP5 | Goat | R&D Systems | AF578 | 1:500 | IF |
| anti-rabbit IgG<br>AlexaFluor 594 | Goat | Life Technologies | A11037 | 1:1000 | IF |
| anti-rabbit IgG<br>AlexaFluor 488 | Donkey | Invitrogen | A-21206 | 1:1000 | IF |
| anti-mouse IgG<br>AlexaFluor 488 | Goat | Life Technologies | A11029 | 1:1000 | IF |
| anti-Goat IgG<br>AlexaFluor 594 | Donkey | Invitrogen | A-11058 | 1:1000 | IF |
| anti-AKT | Rabbit | Cell Signaling Technology | 4691 | 1:1000 | WB |
| anti-Phospho-AKT<br>(ser473) | Rabbit | Cell Signaling Technology | 9271 | 1:1000 | WB |
| anti-Phospho-S6<br>ribosomal protein<br>(Ser235/236) | Mouse | Cell Signaling Technology | 2211S | 1:1000 | WB |
| anti-Beta-actin | Mouse | Sigma-Aldrich | A1978 | 1:2500 | WB |
| Anti-rabbit IgG, HRP-linked | Goat | Cell Signaling Technology | 7074 | 1:5000 | WB |
| Anti-mouse IgG, HRP-linked | Horse | Cell Signaling Technology | 7076 | 1:5000 | WB |

IH, immunohistochemistry; IF, immunofluorescence; WB, Western blot

For immunofluorescence, the developmental stages of nephrons in E15.5 fetal kidneys were stained with antibodies for CYCLIN D1, JAGGED 1, LEF1 and PAX2, alongside with CALBINDIN antibody as an epithelial marker as previously reported^40^. For counting the glomeruli in 2-week kidneys (WT n = 3, *Hsd17b3*^-/-^ n=3), podocytes were stained with PODOCALYXIN antibody. Cell proliferation in the proximal tubule of 2-week kidneys was analyzed by staining phospho-HISTONE H3 and LRP2 as a proximal tubule marker, and expression of IGFBP5 in proximal tubule also confirmed with staining alongside LRP2. Antibodies are listed in Table 2.

Antigen retrieval was performed in a pressure cooker in Tris-HCl EDTA buffer (E18.5 kidneys) or citrate buffer (2-week kidneys). Blocking and primary antibody incubation was carried out as above, followed by a 40-minute incubation with secondary antibodies. Finally, a brief incubation with DAPI was followed by mounting and imaging the slides with Zeiss Axio Imager (Carl Zeiss NTS Ltd.) or scanning them with Pannoramic MIDI fluorescent slide scanner (3DHistech).

### Gene expression analyses and RNA sequencing

Total RNA was extracted from E18.5 kidneys (WT n = 4, *Hsd17b3*^-/-^ n =5) as well as WT E15.5, 0 day and 3 week-old kidneys (n = 3 for all) with Trisure (Bioline) following the manufacturer’s instructions. The RNA integrity of the samples was confirmed with NanoDrop ND-1000 spectrophotometer (Thermo Fisher Scientific). One μg of RNA was treated with DNase Amplification Grade Kit (Thermo Fisher Scientific) and used for cDNA synthesis (SensiFast, Bioline). The cDNA was used to quantify gene expression by qPCR (CFX96 Real-Time PCR detection system, Bio-Rad) with the DyNAmo Flash SYBR Green qPCR Kit (Thermo Fisher Scientific). Gene expression was analysed with primers for mouse *Hnf1b, Hnf4a*, and *Igfbp5*. Data was normalized to the expression of housekeeping genes *L19* and *Ppia*, and expression relative to the WT levels (Hnfs) or lowest expressing time point (Igfbp5) was calculated from Ct values using the ΔΔCt method^41^. Primers are listed Table 3.

**Table 3.**
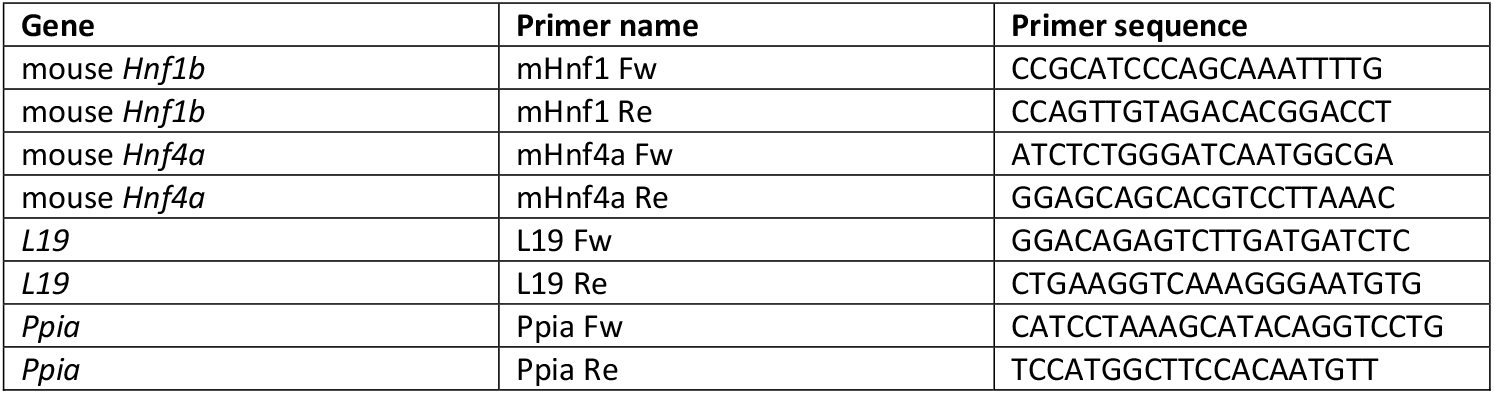
Polymerase chain reaction (PCR) primer sequences.

For RNA sequencing, total RNA was extracted from E16.5 kidneys from the testosterone supplementation experiment as described above. The quality of the RNA samples was confirmed by Bioanalyzer. Library preparation and sequencing were performed by Novogene Co. with Illumina NovaSeq 6000 system. Computational analyses are described separately below.

### Cell culture and proliferation

For proliferation analysis, HEK 293 (ATCC) cells were cultured on poly-L-lysine –coated plates (#P4832, Sigma-Aldrich) and switched to serum-free medium (DMEM/F12, #D2906, Sigma; 1% Penicillin/streptomycin, #A5256701, Gibco; 1% L-Glutamine, #25030-024, Gibco; 20 nM DHT) for 3 hours before changing to fresh serum-free medium with 25, 50 or 100 nM recombinant human IGFBP5 (#875-B5, R&D Systems). Cells were cultured for 24 hours, and the cell proliferation was analyzed with an MTT cell proliferation and cytotoxicity assay kit (#E-CK-A341, Elabscience), measured on an EnSight microplate reader (Perkin Elmer).

### Immunoblotting

For Western blotting, E18.5 KO and WT kidneys were homogenized in RIPA buffer with phosphatase inhibitor, and the proteins separated on Mini-ProteanTGX 4–20% gels (#456-1094, BioRad) and transferred to PVDF membrane with BioRad SemiDry system at 25 V for 30 min. Membranes were blocked in a solution of 3% BSA and 5% fat-free milk. Immunoblotting was performed with antibodies for AKT, Phospho-AKT (pAKT), and β-actin overnight. Secondary staining was performed for 1 h in RT with HRP-conjugated antibodies, and chemiluminescence detection reagents (#NEL122001EA, PerkinElmer) were used for imaging with Fujifilm LAS 4000 gel imager. Signal intensities were quantified with ImageJ software^42^, normalized to β-actin values as loading control. Antibodies are listed in Table 2.

### Computational analyses

The analyses of AR chromatin binding upon testosterone activation in kidney were performed utilizing previously published ChIP-seq data (GSM1146473, GSM1146474, GSM1146475 from GSE47192)^21^ and UCSC Genome Browser (mm9).

RNA-seq data analysis was performed using a combination of programs. The quality of the raw sequencing reads was checked with FastQC tool version 0.11.14^43^. Further analyses were carried out using R version 3.6.1 and Bioconductor version 3.9^44^. The reads were aligned to the UCSC mm10 mouse genome reference, derived from Illumina iGenomes (https://support.illumina.com/sequencing/sequencing_software/igenome.html), using Rsubread package (version 2.0.0)^45^ and its inbuilt Refseq gene annotation. Rsubread was also used to calculate the genewise read counts. Normalization was performed using the calcNormFactors function from the edgeR package version 3.28.0^46^, implementing the trimmed mean of M-values (TMM) normalization method. Statistical testing between sample groups was carried out using ROTS package version 1.14.0^47^, and the differentially expressed genes were selected, requiring a false-discovery rate (FDR) below 0.05 and absolute fold-change (FC) above 1.5.

Gene ontology (GO) enrichment analysis was performed using a threshold-free gene-set enrichment (GSEA) approach with gage package (version 2.36.0)^48^. The range of genes per term required was set between 15 and 250, and the test was performed with comparison scheme ‘as.group’ using the function ‘gs.KSTest’ for the non-parametric Kolmogorov-Smirnov test. The gene lists sorted based on average ranks of both statistical significance and fold-change from the differential expression testing were used as input.

Gene expression data were also analyzed using QIAGEN Ingenuity Pathway Analysis (IPA)^49^. Canonical pathway analysis identified the pathways from QIAGEN IPA library of canonical pathways that were most significant to the data set. Only genes from the dataset that met a P-value cutoff of 0.05 and were associated with a canonical pathway in the QIAGEN Knowledge Base were considered. The significance of the association between the data set and the canonical pathway was measured using a right-tailed Fisher’s Exact Test.

The data set containing gene identifiers and their corresponding expression values was uploaded into the QIAGEN IPA software, and networks were generated. Each identifier was mapped to its corresponding entity molecule in QIAGEN’s Knowledge Base. These molecules were overlaid onto a global molecular network developed from information contained in the QIAGEN’s Knowledge Base. Networks of eligible molecules were then algorithmically generated based on their connectivity, and selected networks were then merged and combined to understand the underlying molecular mechanisms.

### Statistical analyses

Statistical analyses were done using GraphPad Prism 10 software (GraphPad Software). The normality of the data was evaluated based on a Shapiro-Wilk normality test. Statistical difference between two groups was determined by two-tailed Student’s t-test or Mann-Whitney test for normally and non-normally distributed data, respectively. For comparison of multiple groups, one-way ANOVA and Tukey’s multiple comparisons test, or Kruskal-Wallis test and Dunn’s multiple comparisons test were used for normally and non-normally distributed data, respectively. Significance was determined as *P* ≤ 0.05, and results are shown as individual values, with a centerline indicating the mean unless stated otherwise. All experiments presented in the Article were repeated in three independent biological replicates at least. No statistical methods were used to predetermine sample sizes. Data collection and analysis were not performed blinded. No other data points were removed from the experiments.

## Acknowledgements

The authors thank the personnel of Turku Center for Disease Modeling (www.tcdm.fi) supported by the Biocenter Finland, and Medical Bioinformatics Center supported by Biocenter Finland and ELIXIR Finland. We further acknowledge the Histology Core Facility at the Institute of Biomedicine, University of Turku for histological services. This work was supported by Sigrid Juselius Foundation, Jalmari and Rauha Ahokas Foundation, Turku University Foundation, Drug research doctoral programme, University of Turku and Organon RD Finland. The funders had no role in the design, data collection, data analysis, and reporting of this study.

## Author Contributions

Conceptualization, S.K., M.P. and P.S.; formal analysis, A.J., K.R., G.M-N., M.P., A.L. and L.E.; investigation, A.J., H.L., J.A., I.H., and O.M.; resources C.O., S.K., M.P. and P.S.; Writing – Original Draft, A.J. and P.S.; Writing – Review & Editing, all authors; supervision C.O., L.E., S.K., M.P. and P.S.; Funding Acquisition, M.P. and P.S.

## Data availability

The RNA sequencing data has been deposited in the GEO database under accession number GSE280108. All the other data are presented in manuscript or in the supplementary materials. Additional requests can be directed to the corresponding author.

## Supplemental figure legends

**Figure S1**. IPA gene interaction network map. The network was generated by merging the individual networks of the molecules IGFBP5 and mTOR obtained after comparing differential gene expression between WT and *Hsd17b3*-/-. These factors were significantly represented in the top canonical pathway results.

## References

Catterall, J.F., Kontula, K.K., Watson, C.S., Seppänen, P.J., Funkenstein, B., Melanitou, E., Hickok, N.J., Bardin, C.W., and Jänne, O.A. (1986). Regulation of Gene Expression by Androgens in Murine Kidney. In Proceedings of the 1985 Laurentian Hormone Conference Recent Progress in Hormone Research., R. O. Greep, ed. (Academic Press), pp. 71–109. 10.1016/B978-0-12-571142-5.50006-9.

Asadi, F.K., Dimaculangan, D.D., and Berger, F.G. (1994). Androgen regulation of gene expression in primary epithelial cells of the mouse kidney. Endocrinology 134, 1179–1187. 10.1210/endo.134.3.8119157.

Ransick, A., Lindström, N.O., Liu, J., Zhu, Q., Guo, J.J., Alvarado, G.F., Kim, A.D., Black, H.G., Kim, J., and McMahon, A.P. (2019). Single-Cell Profiling Reveals Sex, Lineage, and Regional Diversity in the Mouse Kidney. Dev. Cell 51, 399-413.e7. 10.1016/j.devcel.2019.10.005.

McDonough, A.A., Harris, A.N., Xiong, L. (Ivy), and Layton, A.T. (2023). Sex differences in renal transporters: assessment and functional consequences. Nat. Rev. Nephrol. 20, 21. 10.1038/s41581-023-00757-2.

Veiras, L.C., Girardi, A.C.C., Curry, J., Pei, L., Ralph, D.L., Tran, A., Castelo-Branco, R.C., Pastor-Soler, N., Arranz, C.T., Yu, A.S.L., et al. (2017). Sexual Dimorphic Pattern of Renal Transporters and Electrolyte Homeostasis. J. Am. Soc. Nephrol. JASN 28, 3504–3517. 10.1681/ASN.2017030295.

Harris, A.N., Lee, H.-W., Osis, G., Fang, L., Webster, K.L., Verlander, J.W., and Weiner, I.D. (2018). Differences in renal ammonia metabolism in male and female kidney. Am. J. Physiol. Renal Physiol. 315, F211–F222. 10.1152/ajprenal.00084.2018.

Sabolić, I., Asif, A.R., Budach, W.E., Wanke, C., Bahn, A., and Burckhardt, G. (2007). Gender differences in kidney function. Pflüg. Arch. - Eur. J. Physiol. 455, 397–429. 10.1007/s00424-007-0308-1.

Harris, A.N., Lee, H.-W., Verlander, J.W., and Weiner, I.D. (2020). Testosterone modulates renal ammonia metabolism. Am. J. Physiol. - Ren. Physiol. 318, F922–F935. 10.1152/ajprenal.00560.2019.

Laouari, D., Vergnaud, P., Hirose, T., Zaidan, M., Rabant, M., Nguyen, C., Burtin, M., Legendre, C., Codogno, P., Friedlander, G., et al. (2022). The sexual dimorphism of kidney growth in mice and humans. Kidney Int. 102, 78–95. 10.1016/j.kint.2022.02.027.

Harris, A.N., Castro, R.A., Lee, H.-W., Verlander, J.W., and Weiner, I.D. (2021). Role of the renal androgen receptor in sex differences in ammonia metabolism. Am. J. Physiol. Renal Physiol. 321, F629–F644. 10.1152/ajprenal.00260.2021.

Labrie, F., Luu-The, V., Lin, S.X., Simard, J., and Labrie, C. (2000). Role of 17β-hydroxysteroid dehydrogenases in sex steroid formation in peripheral intracrine tissues. Trends Endocrinol. Metab. 11, 421–427. 10.1016/S1043-2760(00)00342-8.

Sipilä, P., Junnila, A., Hakkarainen, J., Huhtaniemi, R., Mairinoja, L., Zhang, F.P., Strauss, L., Ohlsson, C., Kotaja, N., Huhtaniemi, I., et al. (2020). The lack of HSD17B3 in male mice results in disturbed Leydig cell maturation and endocrine imbalance akin to humans with HSD17B3 deficiency. FASEB J. 34, 6111–6128. 10.1096/fj.201902384R.

O’Shaughnessy, P.J., Baker, P., Sohnius, U., Haavisto, A.M., Charlton, H.M., and Huhtaniemi, I. (1998). Fetal development of Leydig cell activity in the mouse is independent of pituitary gonadotroph function. Endocrinology 139, 1141–1146. 10.1210/endo.139.3.5788.

Silversides, D.W., Price, C.A., and Cooke, G.M. (1995). Effects of short-term exposure to hydroxyflutamide in utero on the development of the reproductive tract in male mice. Can. J. Physiol. Pharmacol. 73, 1582– 1588. 10.1139/y95-718.

Welsh, M., Saunders, P.T.K., Marchetti, N.I., and Sharpe, R.M. (2006). Androgen-Dependent Mechanisms of Wolffian Duct Development and Their Perturbation by Flutamide. Endocrinology 147, 4820–4830. 10.1210/en.2006-0149.

Welsh, M., Saunders, P.T.K., and Sharpe, R.M. (2007). The Critical Time Window for Androgen-Dependent Development of the Wolffian Duct in the Rat. Endocrinology 148, 3185–3195. 10.1210/en.2007-0028.

Welsh, M., Saunders, P.T.K., Fisken, M., Scott, H.M., Hutchison, G.R., Smith, L.B., and Sharpe, R.M. (2008). Identification in rats of a programming window for reproductive tract masculinization, disruption of which leads to hypospadias and cryptorchidism. J. Clin. Invest. 118, 1479–1490. 10.1172/JCI34241.

Junnila, A., Zhang, F.-P., Martínez Nieto, G., Hakkarainen, J., Mäkelä, J.-A., Ohlsson, C., Sipilä, P., and Poutanen, M. (2024). HSD17B1 Compensates for HSD17B3 Deficiency in Fetal Mouse Testis but Not in Adults. Endocrinology 165, bqae056. 10.1210/endocr/bqae056.

Marable, S.S., Chung, E., Adam, M., Potter, S.S., and Park, J.-S. (2018). Hnf4a deletion in the mouse kidney phenocopies Fanconi renotubular syndrome. JCI Insight 3. 10.1172/jci.insight.97497.

Shao, A., Chan, S.C., and Igarashi, P. (2020). Role of Transcription Factor Hepatocyte Nuclear Factor-1β in Polycystic Kidney Disease. Cell. Signal. 71, 109568. 10.1016/j.cellsig.2020.109568.

Pihlajamaa, P., Sahu, B., Lyly, L., Aittomäki, V., Hautaniemi, S., and Jänne, O.A. (2014). Tissue-specific pioneer factors associate with androgen receptor cistromes and transcription programs. EMBO J. 33, 312– 326. 10.1002/embj.201385895.

Adam, M., Potter, A.S., and Potter, S.S. (2017). Psychrophilic proteases dramatically reduce single-cell RNA-seq artifacts: a molecular atlas of kidney development. Dev. Camb. Engl. 144, 3625–3632. 10.1242/dev.151142.

Kim, S., Koppitch, K., Parvez, R.K., Guo, J., Achieng, M., Schnell, J., Lindström, N.O., and McMahon, A.P. (2024). Comparative single-cell analyses identify shared and divergent features of human and mouse kidney development. Dev. Cell 59, 2912-2930.e7. 10.1016/j.devcel.2024.07.013.

Marable, S.S., Chung, E., and Park, J.-S. (2020). Hnf4a Is Required for the Development of Cdh6-Expressing Progenitors into Proximal Tubules in the Mouse Kidney. J. Am. Soc. Nephrol. JASN 31, 2543–2558. 10.1681/ASN.2020020184.

Xiong, L., Liu, J., Han, S.Y., Koppitch, K., Guo, J.-J., Rommelfanger, M., Miao, Z., Gao, F., Hallgrimsdottir, I.B., Pachter, L., et al. (2023). Direct androgen receptor control of sexually dimorphic gene expression in the mammalian kidney. Dev. Cell 58, 2338-2358.e5. 10.1016/j.devcel.2023.08.010.

Duan, C., and Allard, J.B. (2020). Insulin-Like Growth Factor Binding Protein-5 in Physiology and Disease. Front. Endocrinol. 11, 100. 10.3389/fendo.2020.00100.

Gleason, C.E., Ning, Y., Cominski, T.P., Gupta, R., Kaestner, K.H., Pintar, J.E., and Birnbaum, M.J. (2010). Role of Insulin-Like Growth Factor-Binding Protein 5 (IGFBP5) in Organismal and Pancreatic β-Cell Growth. Mol. Endocrinol. 24, 178–192. 10.1210/me.2009-0167.

Xia, Z., Liu, F., Zhang, J., and Liu, L. (2015). Decreased Expression of MiRNA-204-5p Contributes to Glioma Progression and Promotes Glioma Cell Growth, Migration and Invasion. PLOS ONE 10, e0132399. 10.1371/journal.pone.0132399.

Miyakoshi, N., Richman, C., Kasukawa, Y., Linkhart, T.A., Baylink, D.J., and Mohan, S. (2001). Evidence that IGF-binding protein-5 functions as a growth factor. J. Clin. Invest. 107, 73–81. 10.1172/JCI10459.

Chen, X., Yu, Q., Pan, H., Li, P., Wang, X., and Fu, S. (2020). <p>Overexpression of IGFBP5 Enhances Radiosensitivity Through PI3K-AKT Pathway in Prostate Cancer</p>. Cancer Manag. Res. 12, 5409–5418. 10.2147/CMAR.S257701.

Yao, Z., Lin, M., Lin, T., Gong, X., Qin, P., Li, H., Kang, T., Ye, J., Zhu, Y., Hong, Q., et al. (2022). The expression of IGFBP-5 in the reproductive axis and effect on the onset of puberty in female rats. Reprod. Biol. Endocrinol. 20, 100. 10.1186/s12958-022-00966-7.

Volovelsky, O., Nguyen, T., Jarmas, A.E., Combes, A.N., Wilson, S.B., Little, M.H., Witte, D.P., Brunskill, E.W., and Kopan, R. (2018). Hamartin regulates cessation of mouse nephrogenesis independently of Mtor. Proc. Natl. Acad. Sci. 115, 5998–6003. 10.1073/pnas.1712955115.

Mendoza, M.C., Er, E.E., and Blenis, J. (2011). The Ras-ERK and PI3K-mTOR pathways: cross-talk and compensation. Trends Biochem. Sci. 36, 320–328. 10.1016/j.tibs.2011.03.006.

Mendonca, B.B., Gomes, N.L., Costa, E.M.F., Inacio, M., Martin, R.M., Nishi, M.Y., Carvalho, F.M., Tibor, F.D., and Domenice, S. (2017). 46,XY disorder of sex development (DSD) due to 17β-hydroxysteroid dehydrogenase type 3 deficiency. J. Steroid Biochem. Mol. Biol. 165, 79–85. 10.1016/j.jsbmb.2016.05.002.

van den Driesche, S., Kilcoyne, K.R., Wagner, I., Rebourcet, D., Boyle, A., Mitchell, R., McKinnell, C., Macpherson, S., Donat, R., Shukla, C.J., et al. (2017). Experimentally induced testicular dysgenesis syndrome originates in the masculinization programming window. JCI Insight 2, e91204. 10.1172/jci.insight.91204.

Abitbol, C.L., and Rodriguez, M.M. (2012). The long-term renal and cardiovascular consequences of prematurity. Nat. Rev. Nephrol. 8, 265–274. 10.1038/nrneph.2012.38.

Borrelli, S., Provenzano, M., Gagliardi, I., Ashour, M., Liberti, M.E., De Nicola, L., Conte, G., Garofalo, C., and Andreucci, M. (2020). Sodium Intake and Chronic Kidney Disease. Int. J. Mol. Sci. 21, 4744. 10.3390/ijms21134744.

Nanra, R.S., Stuart-Taylor, J., de Leon, A.H., and White, K.H. (1978). Analgesic nephropathy: etiology, clinical syndrome, and clinicopathologic correlations in Australia. Kidney Int. 13, 79–92. 10.1038/ki.1978.11.

Nilsson, M.E., Vandenput, L., Tivesten, Å., Norlén, A.K., Lagerquist, M.K., Windahl, S.H., Börjesson, A.E., Farman, H.H., Poutanen, M., Benrick, A., et al. (2015). Measurement of a comprehensive sex steroid profile in rodent serum by high-sensitive gas chromatography-tandem mass spectrometry. Endocrinology 156, 2492–2502. 10.1210/en.2014-1890.

Ihermann-Hella, A., Lume, M., Miinalainen, I.J., Pirttiniemi, A., Gui, Y., Peränen, J., Charron, J., Saarma, M., Costantini, F., and Kuure, S. (2014). Mitogen-Activated Protein Kinase (MAPK) Pathway Regulates Branching by Remodeling Epithelial Cell Adhesion. PLoS Genet. 10. 10.1371/journal.pgen.1004193.

Livak, K.J., and Schmittgen, T.D. (2001). Analysis of Relative Gene Expression Data Using Real-Time Quantitative PCR and the 2−ΔΔCT Method. Methods 25, 402–408. 10.1006/meth.2001.1262.

Schneider, C.A., Rasband, W.S., and Eliceiri, K.W. (2012). NIH Image to ImageJ: 25 years of image analysis. Nat. Methods 9, 671–675. 10.1038/nmeth.2089.

Babraham Bioinformatics - FastQC A Quality Control tool for High Throughput Sequence Data https://www.bioinformatics.babraham.ac.uk/projects/fastqc/.

Huber, W., Carey, V.J., Gentleman, R., Anders, S., Carlson, M., Carvalho, B.S., Bravo, H.C., Davis, S., Gatto, L., Girke, T., et al. (2015). Orchestrating high-throughput genomic analysis with Bioconductor. Nat. Methods 12, 115–121. 10.1038/nmeth.3252.

Liao, Y., Smyth, G.K., and Shi, W. (2019). The R package Rsubread is easier, faster, cheaper and better for alignment and quantification of RNA sequencing reads. Nucleic Acids Res. 47, e47. 10.1093/nar/gkz114.

McCarthy, D.J., Chen, Y., and Smyth, G.K. (2012). Differential expression analysis of multifactor RNA-Seq experiments with respect to biological variation. Nucleic Acids Res. 40, 4288–4297. 10.1093/nar/gks042.

Suomi, T., Seyednasrollah, F., Jaakkola, M.K., Faux, T., and Elo, L.L. (2017). ROTS: An R package for reproducibility-optimized statistical testing. PLOS Comput. Biol. 13, e1005562. 10.1371/journal.pcbi.1005562.

gage Bioconductor. http://bioconductor.org/packages/gage/.

Krämer, A., Green, J., Pollard, J., and Tugendreich, S. (2014). Causal analysis approaches in Ingenuity Pathway Analysis. Bioinforma. Oxf. Engl. 30, 523–530. 10.1093/bioinformatics/btt703.

