## Supplemental Figure 1 for "Testosterone exposure during fetal masculinization programming window determines the kidney size in adult mice"

**Supplementary figure 1**

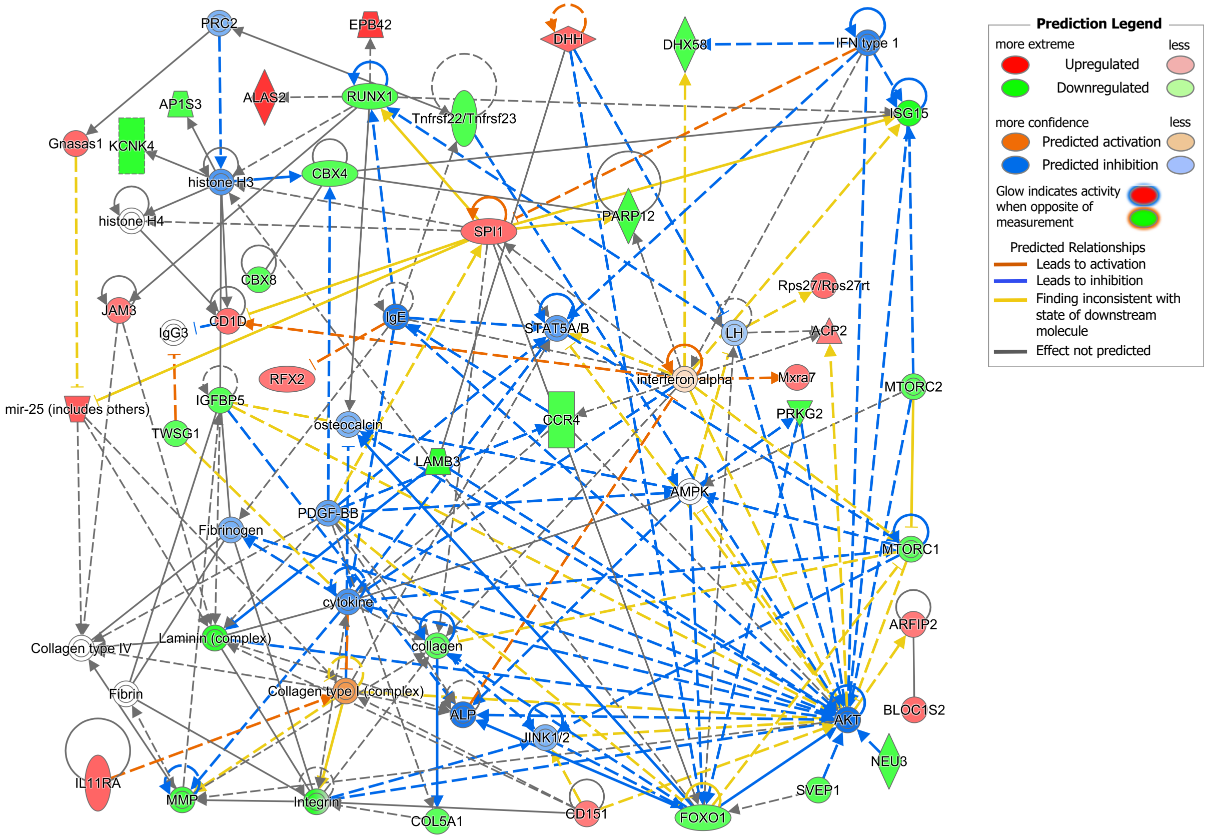


© 2000-2024 QIAGEN. All rights reserved.

**Supplementary figure 1.** IPA gene interaction network map. The network was generated by merging the individual networks of the molecules IGFBP5 and mTOR obtained after comparing differential gene expression between WT and Hsd17b3-/-. These factors were significantly represented in the top canonical pathway results.
